# Genetic Disruption at the CIP2A Locus Modulates T Cell Responses and Attenuates Experimental Autoimmune Encephalomyelitis

**DOI:** 10.64898/2026.08.28.746989

**Authors:** Mohd Moin Khan, Inna Starskaia, Emilie Rydgren, Sini Junttila, Johannes Smolander, Meraj Hasan Khan, Rahul Biradar, Roosa Kattelus, Anne Ahtikoski, Maria Gardberg, Mudassir Meraj Banday, Uzma Riyaz, Joonas Khabbal, Emrah Yatkin, Li Tian, Peggy P. Ho, Tapio Lönnberg, Laura L. Elo, Lawrence Steinman, Jukka Westermarck, Omid Rasool, Riitta Lahesmaa, Ubaid Ullah Kalim

**Affiliations:** Turku Bioscience Centre, University of Turku and Åbo Akademi University, Turku, Finland; Brigham and Women’s Hospital, Harvard Medical School, Boston, Massachusetts, US; InFLAMES Research Flagship Center, University of Turku, Turku, Finland; Depratment of Histology, Turku University Central Hospital (TYKS) Turku FI-20520, Finland; Institute of Biomedicine, University of Turku, Turku, Finland; Central Animal Laboratory, University of Turku, Turku FI-20520, Finland; Department of Psychology, Faculty of Medicine, University of Helsinki; Department of Neurology and Neurological Sciences, Stanford University School of Medicine, Beckman Center for Molecular Medicine, Stanford, CA 94305; Institute for Immunity, Transplantation, and Infection, Stanford University, Stanford, CA 94305

## Abstract

Multiple sclerosis (MS) is a chronic autoimmune disease of the central nervous system (CNS) driven by pathogenic T cell-mediated inflammation. Fingolimod (FTY720), an approved therapy for MS, is an established activator of protein phosphatase 2A (PP2A). However the contribution of PP2A in autoimmune neuroinflammation remains incompletely understood. Here, we addressed this question using experimental autoimmune encephalomyelitis (EAE), a murine model of MS, in mice carrying a genetic disruption of the locus encoding cancerous inhibitor of protein phosphatase 2A (CIP2A), an endogenous inhibitor of PP2A.

Mice with disruption of the CIP2A locus, the knock out (KO) mice, exhibited attenuated EAE severity compared with wild-type (WT) controls. Histological and flow-cytometric analyses revealed markedly reduced infiltration of mononuclear cells, including CD4^+^ and CD4^+^CXCR6^+^ encephalitogenic T cells, in the CNS of diseased KO mice. Reduced numbers of these T cell populations were also observed in peripheral lymphoid organs of the Cip2a-deficient mice during EAE, while T cell abundance was comparable under steady-state conditions, suggesting impaired activation-induced expansion rather than altered homeostasis or migration.

Single-cell RNA sequencing of CNS and lymph node immune cells revealed changes in cell-type abundance and gene expression. Notably, Il17a expression was reduced in CNS CD8+ T cells and showed a similar trend in γδ T cells. Together, our findings reveal that genetic disruption at the CIP2A locus attenuates EAE, possibly by limiting the expansion and accumulation of encephalitogenic T cell populations in CNS. These results identify the CIP2A locus as a previously unrecognized regulator of T cell-driven autoimmune neuroinflammation and provide new insights into mechanisms that restrain pathogenic T cell responses during EAE.

## INTRODUCTION

Multiple sclerosis (MS) is a chronic autoimmune disease of the central nervous system (CNS) characterized by immune-mediated demyelination and neurodegeneration. Experimental autoimmune encephalomyelitis (EAE) is a widely used murine model that recapitulates key pathological and immunological features of MS, including the accumulation of autoreactive T cells in the CNS and subsequent inflammation-driven tissue damage. Among immune cell populations, CD4+ T cells—particularly Th17 cells—play a central role in the initiation and progression of CNS autoimmunity by producing pro-inflammatory cytokines such as interleukin-17A (IL-17A) and by orchestrating recruitment and activation of additional immune cells (*1*). Additionally, other T cell subsets—including IL-17–producing CD8+ T cells (Tc17 cells) and γδ T cells—have been implicated in the pathology of EAE and MS (*2*–*4*). Understanding the molecular regulators that control the activation, expansion, and effector functions of these cells is therefore critical for identifying new therapeutic targets.

Fingolimod (FTY720), an approved therapy of MS is a well-established activator of protein phosphatase 2A (PP2A) activity by blocking the direct protein interaction between PP2A and its inhibitor protein SET (*5*). In tissues, the majority of FTY720 remains unphosphorylated and this form of the drug selectively activates PP2A (*6*). Therefore, the role of PP2A in attenuating MS activity might have been underappreciated. In addition to SET, cancerous inhibitor of protein phosphatase 2A (CIP2A) is another endogenous inhibitor of PP2A. Although CIP2A has been extensively studied in cancer biology (*7, 8*), its role in immune regulation and autoimmune disease remains incompletely understood.

Work from our group demonstrated that CIP2A regulates T cell activation and Th17 cell differentiation in human and mouse T cells (*9*). However, whether CIP2A contributes to autoimmune pathology in vivo, and the mechanisms through which it may influence neuroinflammation, remain unknown.

To address this question, we induced EAE in mice carrying a gene-trap disruption of Cip2a (hereafter referred to as knockout [KO] mice) and characterized disease development using clinical assessment, histological and flow-cytometric analyses, and single-cell RNA sequencing of immune cells isolated from the CNS and peripheral lymphoid organs. These studies identify genetic disruption at the CIP2A locus as a modulator of autoimmune neuroinflammation and reveal cellular and transcriptional changes associated with attenuated EAE.

## RESULTS

### Genetic disruption at the CIP2A locus attenuates EAE in mice

We earlier reported that CIP2A-silencing leads to increased Th17 cell differentiation in both human and mouse T cells (*9*). Based on this observation, we hypothesized that CIP2A deficiency would exacerbate disease severity in EAE, where Th17 cells are key mediators of disease progression. To investigate the function of CIP2A in CNS autoimmunity, we utilized CIP2A^HOZ^ mice, referred to as knock out (KO), where *Cip2a* gene is disrupted by the insertion of the pGT0Lxf vector carrying a β-geo fusion reporter gene in the intron 1. This genetic modification results in the expression of a truncated CIP2A protein consisting of exon 1 fused to β-geo. As previously reported (*10*), KO mice were viable, fertile, and devoid of any obvious morphological abnormalities. Further, we did not observe gross defects in the size of the spleen, lymph nodes and thymus of KO mice (**Fig. 1A**).

**Figure 1.**
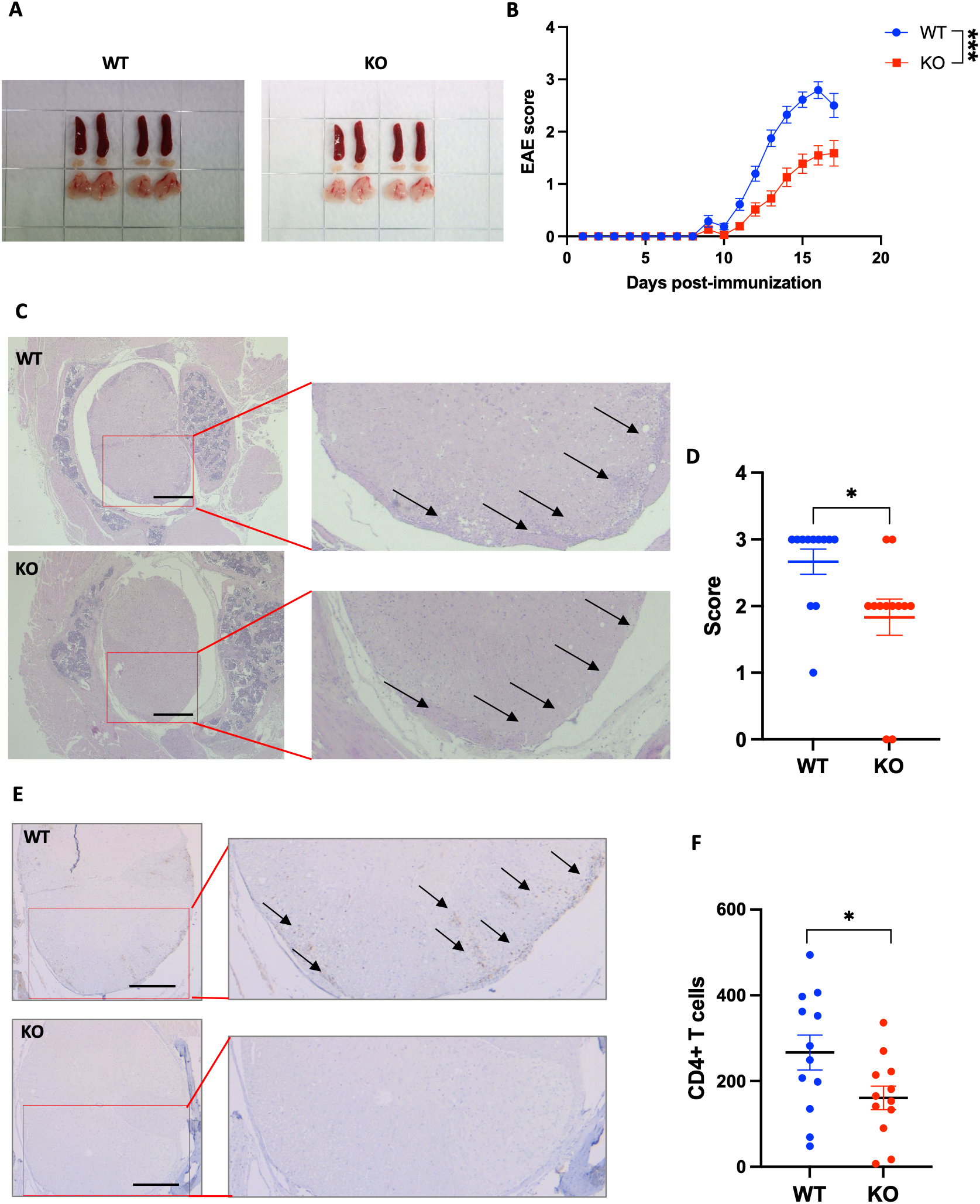
Genetic disruption at the Cip2a locus ameliorates EAE and reduces CNS immune cell infiltration. **(A)** Spleen and lymph nodes from WT (left) and KO (right) animals. **(B)** Clinical EAE scores in WT (N=40) and KO (N=31) mice. Statistical significance for data in panel B was assessed using a linear mixed-effects model in GraphPad Prism, with days post-immunization, genotype, and their interaction included as fixed effects. *** denote P value <0.0001, respectively. Error bars denote standard error of mean (SEM). **(C)** Representative hematoxylin and eosin (H&E)–stained spinal cord sections from WT and KO mice at peak of EAE severity. **(D)** Quantification of inflammatory cell infiltration in spinal cord sections scored from H&E staining (mean+/-SEM). **(E)** Representative immunohistochemical staining of CD4+ T cells in spinal cord sections from WT and KO mice with EAE. **(F)** Quantification of CD4+ T cell numbers in spinal cord sections (mean+/-SEM). Each dot represents an individual mouse. Statistical significance unless specified was determined using unpaired two-tailed Student’s t-test, where *, ** represent p<0.05, 0.01, respectively.

EAE was induced by immunizing KO and WT mice with MOG_35-55_ peptides in complete Freund’s adjuvant, followed by administration of pertussis toxin (see Methods for details). Interestingly, mice that were lacking the *Cip2a* gene showed significantly ameliorated disease in KO compared to WT (**Fig. 1B**), and the effect was more pronounced in females (**Fig. S1**).

### KO mice had reduced CNS infiltration of mononuclear cells and T cells during EAE

To assess the impact of the genetic disruption at Cip2a locus on the immune cell infiltration during EAE, we performed histological analyses of the brain and spinal cord at the peak of disease. WT and KO mice were sacrificed, and spinal cord sections were examined by hematoxylin and eosin (H&E) staining. Histopathological evaluation revealed a significant reduction in the number of infiltrating mononuclear cells (MNCs) in the spinal cords of KO mice compared with WT controls (**Fig. 1C-D**). Immunohistochemical analysis further demonstrated a marked decrease in CD4+ T cells in the spinal cords of KO mice (**Fig. 1E-F**). Infiltrating cells were predominantly localized within perivascular spaces and the parenchyma.

Consistent with these histological findings, flow cytometric analysis of CNS-isolated cells from EAE-diseased mice revealed significantly reduced frequencies and numbers of CD4+ T cells (**Fig. 2A-B**), as well as CD4+ CXCR6+ encephalitogenic Th17 cells (*11*) (**Fig. 2C-D**), in the CNS of KO mice.

**Figure 2.**
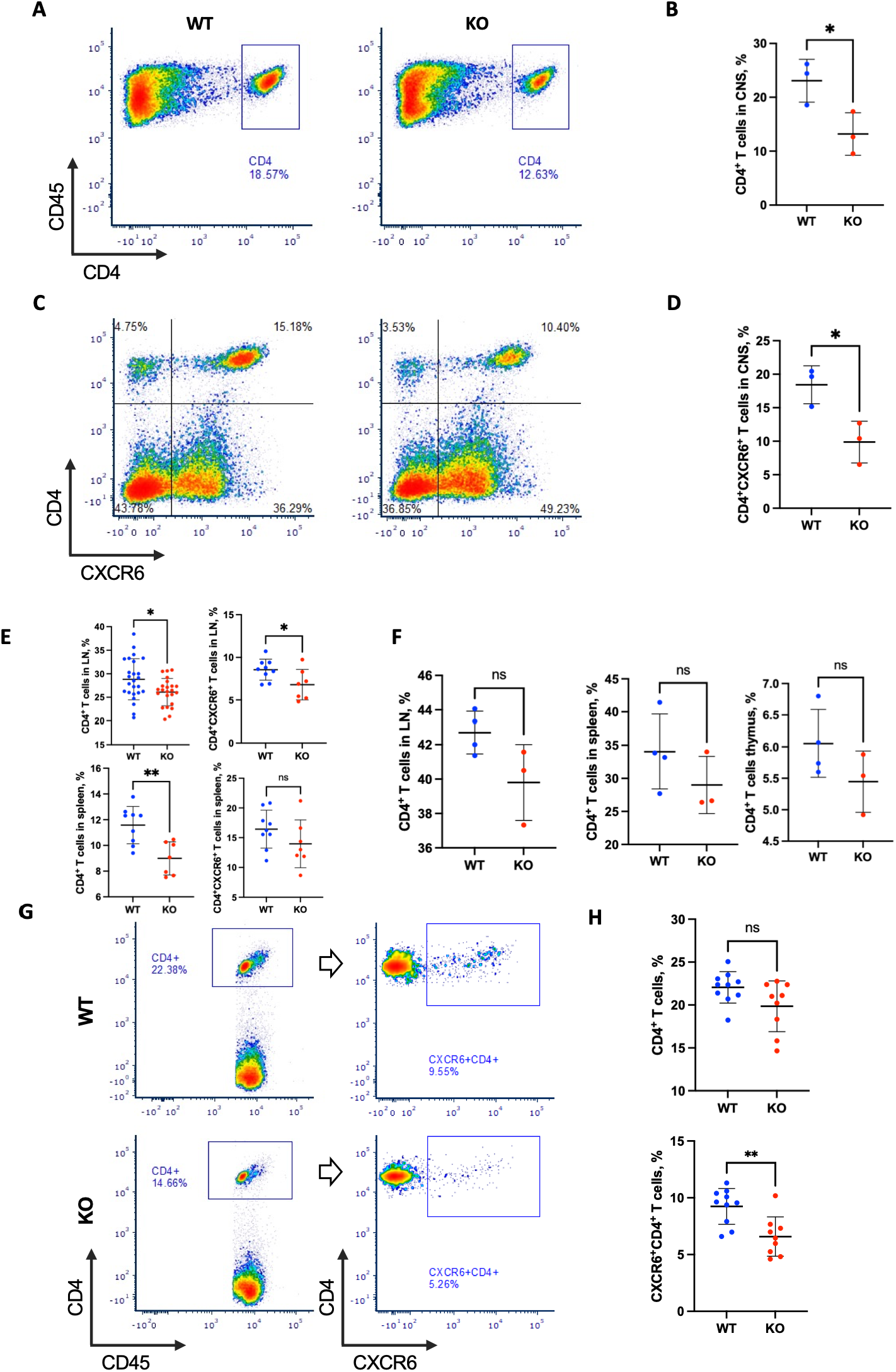
Reduced CD4^+^ T cells in the CNS and periphery of Cip2a-deficient mice with EAE. **(A–B)** Flow-cytometric analysis of CD4^+^ T cells within CD45^+^ mononuclear cells (MNCs) isolated from the CNS of female wild-type (WT) and Cip2a-deficient homozygous (KO/HOZ) mice at the peak of EAE. **(C–D)** Flow-cytometric detection of CD4^+^CXCR6^+^ T cells within CD45^+^ CNS-infiltrating MNCs from WT and KO mice with EAE. **(E)** Frequencies of CD4^+^ and CD4^+^CXCR6^+^ T cells in lymph node and spleen cells from WT and KO mice at the peak of EAE severity. Each dot represents an individual mouse. **(F)** Frequencies of CD4^+^ T cells in lymph nodes and spleens of WT and KO mice under healthy (non-EAE) conditions. **(G-H)** Frequencies of CD4^+^ and CD4^+^CXCR6^+^ T cells from WT and KO splenocytes after 72 h in vitro restimulation with MOG^35–55^ peptide. Statistical significance was determined using unpaired two-tailed Student’s t-test, where *, **, represent p<0.05, 0.01, respectively. ns denotes not significant.

We then asked if the reduced cell number in CNS was due to reduced migration or due to reduced proliferation of cells in KO mice. Interestingly, we observed reduced numbers of CD4+ T cells and CD4+ CXCR6+ T cells in the draining lymph nodes and spleens of KO mice at the peak of EAE (**Fig. 2E-F**). This suggests that the diminished accumulation of these cells in the CNS is unlikely to be solely due to impaired migration and may instead reflect defects in the expansion of encephalitogenic T cells during disease, and thereby limiting activation-induced clonal expansion and survival of encephalitogenic T cells.

To determine whether these differences were attributable to baseline alterations in T cell homeostasis, we analyzed CD4+ T cell populations in the spleen, lymph nodes, and thymus of KO and WT mice without EAE induction. In agreement with earlier observations (*10, 12*), no significant differences in CD4+ T cell numbers were observed between genotypes under steady-state conditions (**Fig. 2F**), suggesting that activation-induced T cell responses rather than homeostatic T cell development or maintenance are affected in KO mice.

Lymphocytes isolated from the spleen and lymph nodes of WT and KO mice with EAE were restimulated in vitro with MOG^35–55^ peptide for 72 h. Cultures from KO mice exhibited significantly reduced frequencies of CD4+ CXCR6+ T cells compared with WT controls (**Fig. 2G-H**). These results suggest that the expansion of MOG^35–55^ specific T cells during EAE is limited in KO mice, possibly contributing to the attenuated disease observed in KO mice.

### Single-cell RNA-seq analysis of immune cells in the CNS and lymph nodes during EAE reveals transcriptional differences in KO mice

To further characterize immune responses in KO mice, we performed single-cell RNA sequencing (scRNA-seq) on mononuclear cells (MNCs) isolated from the brain and spinal cord of WT and KO mice at the peak of EAE. CNS MNCs were enriched using a Percoll gradient and processed for scRNA-seq. To account for biological and technical variability, we profiled MNC populations from two independent pools per genotype, each pool containing cells from four mice with EAE.

Unsupervised clustering identified 21 distinct cell clusters (**Fig. 3A**), which were annotated based on established marker gene expression. Overall, cell-type abundances were comparable between WT and KO mice, with no significant differences in the proportions of major immune populations (**Table S1**). Despite similar cellular composition, differential gene expression analysis identified 18 genes that were significantly differentially expressed (DE) between WT and KO mice (false discovery rate [FDR] < 0.05; fold change ≥ 1.5) in one or more clusters (**Fig. 3B**). Differentially expressed (DE) genes was detected within several cell types, including neutrophils, B cells and CD4 T cells (**Table S2**).

**Figure 3.**
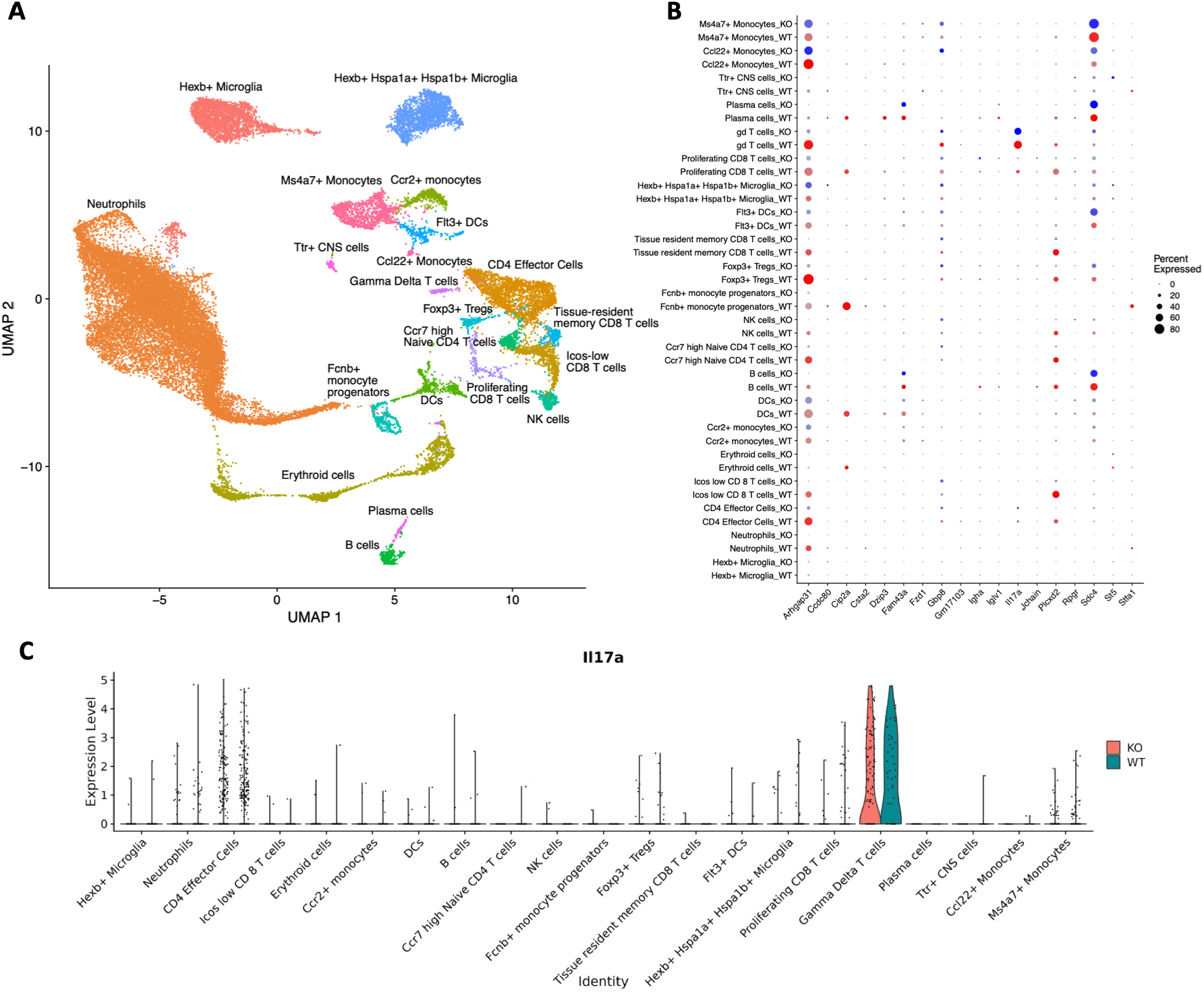
scRNA-seq analysis of CNS mononuclear cells from WT and KO mice at peak of EAE severity. (**A**) UMAP visualization of CNS mononuclear cells (MNCs) isolated from WT and KO mice at peak of EAE severity, showing the major cellular clusters identified by unsupervised clustering. (**B**) Dot plot displaying DE genes (FDR < 0.05; fold change ≥ 1.5) between WT and KO mice across clusters. Dot size indicates the proportion of cells expressing each gene; red and blue reflect higher expression in WT and KO, respectively. (**C**) Violin plot illustrating Il17a expression across immune cell clusters in WT and KO mice.

Interestingly, we observed reduced expression of Il17a in CD8+ T cells from KO mice compared to WT controls (**Fig. 3B**). A similar trend toward reduced Il17a expression was observed in γδ T cells from KO mice; however, this difference did not reach statistical significance. Il17a expression was detected predominantly in γδ T cells, and to a lesser extent in CD4+ and CD8+ T cells (**Fig. 3C**).

In addition to CNS-resident immune cells, we performed scRNA-seq analysis of draining lymph node cells from five WT and five KO mice with EAE. Clustering analysis identified several clusters (**Fig. 4A**). Interestingly, our findings indicated not only variations in gene expression profiles but also differences in the proportions of various cell types. Th17 cells, T follicular helper (Tfh) cells, and plasma cells were significantly less abundant in the KO mice compared to WT mice **(Fig. 4B**). Differential expression analysis identified 36 genes that were significantly altered between WT and KO mice (FDR < 0.05; fold change ≥ 1.5) in one or more lymph node cell clusters (**Fig. 4C, Table S3**). Notably, six genes (Cip2a, Dzip3, Arhgap31, Plcxd2, Csta2, and Stfa1) were downregulated in KO mice in both CNS and lymph node datasets, suggesting partially overlapping transcriptional programs affected in KO mice across lymph node and CNS immune compartments.

**Figure 4.**
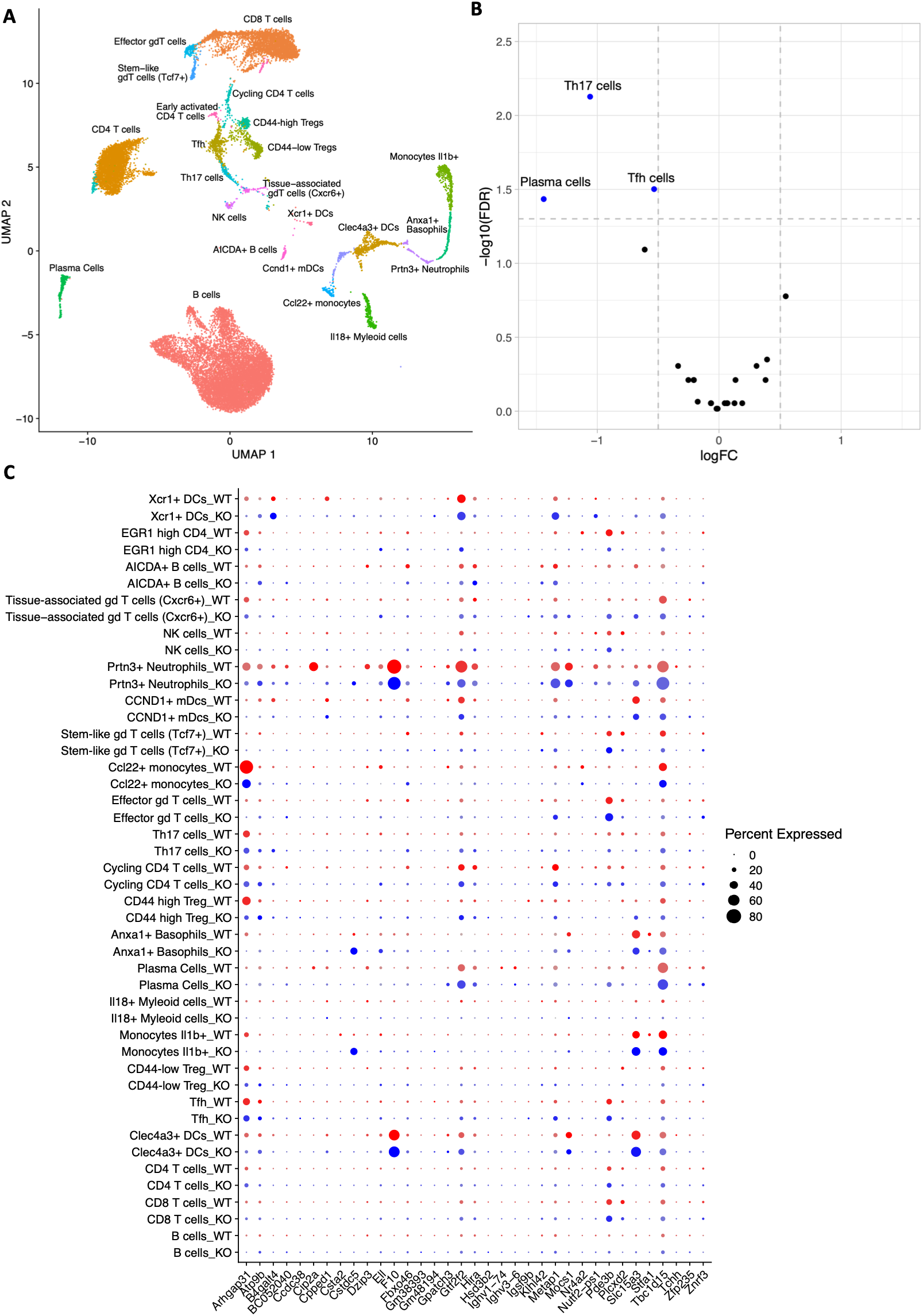
scRNA-seq analysis of cells from the lymph node from WT and KO mice at peak EAE. (**A**) UMAP visualization of lymph node cells isolated from WT and Cip2a-deficient (KO) mice at peak EAE, showing major cellular clusters identified by unsupervised clustering. (**B**) Volcano plots showing the differential abundance analysis between WT and KO mice. Significantly different cell types (edgeR: FDR<0.05, LogFC> |0.5|) are shown on the plot. (**C**) Dot plot displaying DE genes (FDR < 0.05; fold-change ≥ 1.5) between WT and KO mice across clusters. Dot size indicates the proportion of cells expressing each gene; red and blue reflect higher expression in WT and KO, respectively.

### Insertion of the β-geo cassette at the Cip2a locus also disrupts Dzip3 expression

To investigate the overlap between DE genes identified in both the CNS and lymph node scRNA-seq datasets, we examined the six genes that were consistently downregulated in KO mice across both tissues. Notably, among these downregulated genes was Dzip3, which is located about 50 base pairs upstream of Cip2a, with the former transcribed on the reverse strand, whereas the latter is transcribed on the forward strand (**Fig. 5A**). Consistent with their close genomic location, they were also coregulated as identified by the GeneNetwork Assisted Diagnostic Optimization analysis (*13*) exploiting 31,499 public human RNA-seq samples (**Fig. S2**).

**Figure 5.**
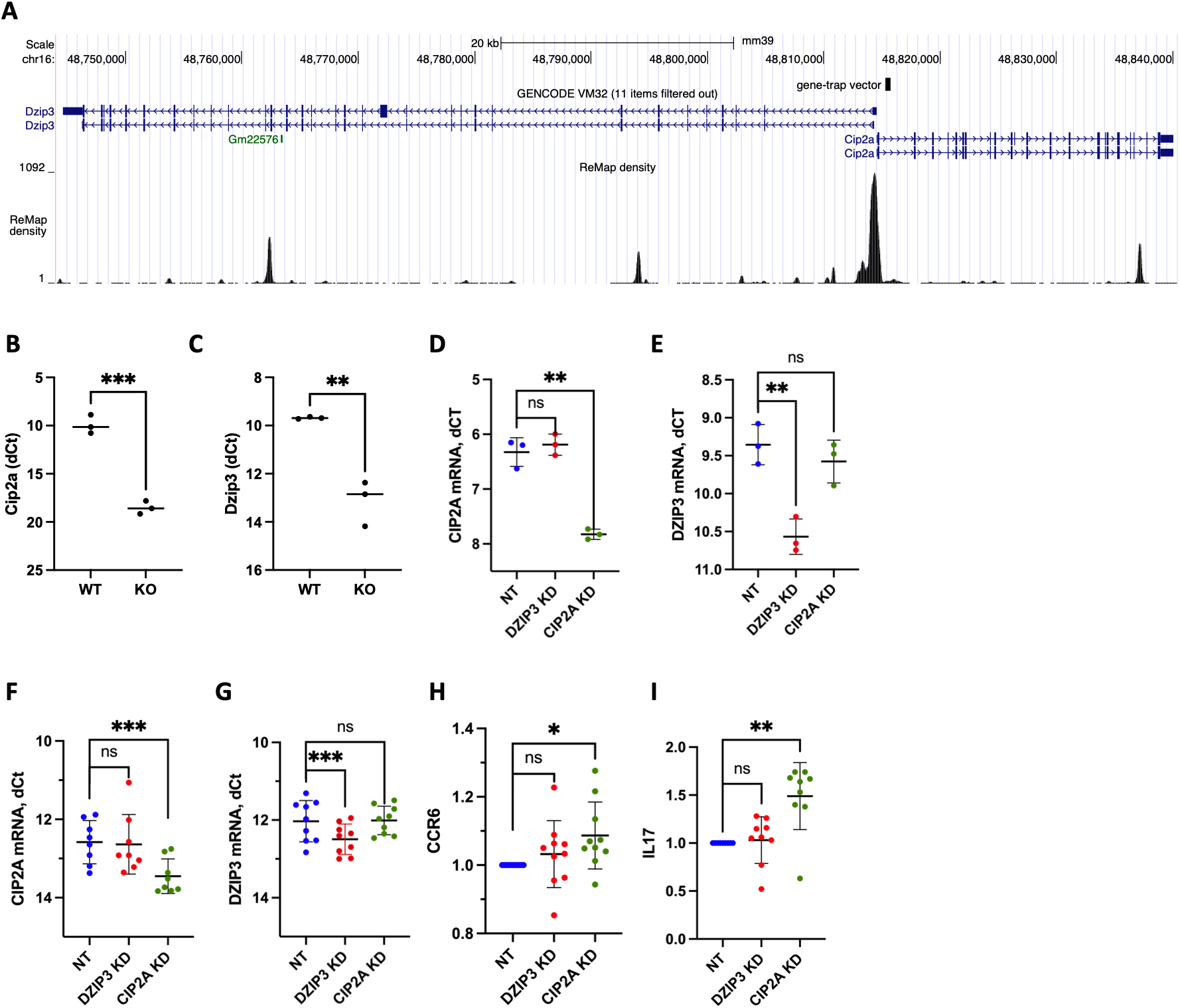
CIP2A and DZIP3 do not regulate each other and CIP2A-but not DZIP3-silencing upregulates IL17A and CCR6. (**A**) UCSC browser shot showing the location of Cip2a and Dzip3 in the mouse genome. Site of gene-trap vector insertion are shown. (**B-C**) Expression of Cip2a and Dzip3 in WT and KO mice. (**D-E**) The effect of siRNA mediated silencing of DZIP3 and CIP2A on their expression in mouse Th17 cells. (**F-G**) The effect of siRNA mediated silencing of DZIP3 and CIP2A on their expression in human Th17 cells. (**H-I**) The effect of siRNA mediated silencing of DZIP3 and CIP2A on the surface expression of CCR6 and secretory IL-17A by in vitro differentiated human Th17 cells at 72 hours. Statistical significance was determined using unpaired two-tailed Student’s t-test, where *, **, and ***, represent p<0.05, 0.01, 0.001, respectively. ns denotes not significant.

This genomic arrangement raised the possibility that insertion of the β-geo reporter cassette into the first intron of Cip2a may have disrupted Dzip3 transcription, potentially by interfering with its promoter region, given the transcription start site of Dzip3 being approximately 750 bp from the cassette insertion site.

Consistent with this possibility, reduced Dzip3 expression in KO mice was not restricted to animals with EAE, but was also observed under steady-state conditions. CD4+ T cells isolated from the spleens of naive KO mice exhibited significantly lower levels of Dzip3 expression compared with WT controls (**Fig. 5B-C**), indicating that Dzip3 downregulation occurs independently of neuroinflammation and is present prior to EAE induction. These observations suggested two potential mechanisms: (i) Cip2a regulates Dzip3 expression, or (ii) the cassette insertion directly disrupts Dzip3 transcription.

We previously showed that silencing of CIP2A in human Th17 cells does not alter DZIP3 expression (*9*), arguing against a direct regulatory relationship. However, to exclude a potential species-specific effect, we tested whether Cip2a regulates Dzip3 expression in mouse Th17 cells. To this end, murine CD4+ T cells were transfected with Cip2a-specific siRNAs, differentiated under Th17-polarizing conditions, and analyzed for Dzip3 expression. Although Cip2a expression was efficiently silenced, Dzip3 expression remained unchanged (**Fig. 5D-E**), indicating that Cip2a does not regulate Dzip3 transcription in mouse Th17 cells. Together, these results suggest that the β-geo cassette insertion disrupts Dzip3 expression independently of Cip2a.

Given that we previously observed increased IL-17A and CCR6 expression following CIP2A silencing in human and mouse Th17 cells (*9*), these findings raised the possibility that the observed Th17 phenotypes could potentially be attributable to Dzip3 deficiency rather than Cip2a loss. To address this, we independently silenced CIP2A or DZIP3 in human Th17 cells and assessed gene expression using TaqMan assays, as well as IL-17A secretion and CCR6 surface expression by ELISA and flow cytometry, respectively. The siRNAs were highly specific, and neither CIP2A nor DZIP3 knockdown altered the expression of the other gene (**Fig. 5F-G**). Importantly, while CIP2A silencing resulted in increased IL-17A secretion and CCR6 expression, silencing of DZIP3 had no effect on IL-17A secretion or CCR6 surface expression (**Fig. 5H-I**).

These data confirm that the previously reported effects of CIP2A loss on IL-17A and CCR6 expression in human and mouse Th17 cells (*9*) are attributable to CIP2A deficiency and are not secondary to altered DZIP3 expression. However, given the disruption of Dzip3 in KO mice, it remains to be determined whether the protection from EAE observed in these animals is mediated predominantly by loss of Cip2a, loss of Dzip3, or a combined effect of both genes.

## DISCUSSION

Fingolimod (FTY720) is a clinically approved therapy for MS and is thought to act primarily by sequestering autoreactive lymphocytes in lymph nodes, thereby preventing their infiltration into the central nervous system (*14*). Notably, FTY720 has also been shown to activate PP2A (*15*), raising the possibility that at least part of its therapeutic effect in EAE and MS may involve PP2A activation (*16*). However, genetic evidence for the role of PP2A inhibitor proteins in EAE susceptibility has been missing. In the present study, histological and flow cytometric analyses demonstrated a marked reduction in mononuclear cell infiltration in the CNS of KO mice during EAE, particularly among CD4+ T cells. Reduced numbers of CD4+ T cells were also observed in peripheral lymphoid organs at peak disease, whereas baseline T cell numbers were comparable between genotypes under steady-state conditions. These findings suggest that in KO mice homeostatic T cell development is not impaired, but rather the expansion or persistence of pathogenic T cell populations is altered during inflammatory responses.

Single-cell RNA sequencing further revealed that in KO mice transcriptional programs rather than CNS immune cell composition are affected. In the CNS, Il17a expression was reduced in CD8+ T cells of KO mice and showed a similar trend in γδ T cells. In lymph nodes of KO mice, we found reduced abundance of Th17 cells, T follicular helper cells, plasma cells, and CD44hi regulatory T cells. Together, these findings indicate that disruption of CIP2A locus inhibits inflammatory gene expression and activation-induced immune cell expansion during EAE.

In addition to effects on gene expression in CNS, KO mice had reduced infiltration of CXCR6+ T cells, a population implicated in the generation of highly pathogenic Th17 cells during neuroinflammation (*11*). Recent studies suggest that IL-17-producing stem-like T cells serve as a reservoir for encephalitogenic Th17 cells, highlighting how reduced availability of CXCR6+ T cells may limit disease severity.

CIP2A has previously been shown to negatively regulate IL-17A expression during Th17 differentiation in T cells (*9*). Given the pathogenic role of Th17 cells in EAE, one would expect that Cip2a deficiency exacerbates disease severity. However, we observed attenuated disease in the KO mice. The apparent contradiction between enhanced Th17 polarization observed upon CIP2A loss in vitro and reduced disease severity in vivo in the KO mice may reflect the functional heterogeneity of Th17 cells. While IL-17A is often associated with pathogenicity, IFN-γ–producing Th17 cells have been identified as particularly encephalitogenic (*17, 18*). We previously showed that in human CIP2A deficiency increases IL-17A while reducing IFN-γ production (*9*), suggesting that loss of CIP2A may skew Th17 cells toward a less pathogenic phenotype in vivo.

Our scRNA-seq analyses also revealed downregulation of Dzip3 in the cells from KO mice across CNS and lymph node compartments. Dzip3 (DAZ Interacting Zinc Finger Protein 3) is an E3 Ubiquitin ligase which also acts as a transcriptional coactivator of Estrogen Receptor alpha (*19*). We demonstrate that Dzip3 downregulation results from disruption at the Cip2a locus rather than transcriptional regulation by CIP2A, but the functional studies in human Th17 cells exclude a role for DZIP3 in regulating IL-17A or CCR6. Given the genomic disruption and the known role of estrogen receptor signaling in EAE and MS (*20*–*22*), the impact of direct Dzip3 inhibition in EAE development should be confirmed in future studies.

Should the observations reported here be mediated through CIP2A, our results provide support for the therapeutic role for PP2A reactivation in preventing MS development. Considering the lack of any detrimental developmental or physiological effects in Cip2a-deficient mice (*8, 10, 12*), and decades of experience in using PP2A reactivating therapy fingolimod in MS patients, direct pharmacological CIP2A inhibition could provide an interesting future alternative therapeutic approach for MS.

## MATERIALS AND METHODS

### Animals

Cip2a gene-deficient mice (*10*) and C57BL/6J control male and female mice, aged 8-12 weeks, were used for EAE studies. These mice were bred in a pathogen-free animal facility at the University of Turku. All animal protocols were approved by the Central Animal Laboratory, University of Turku Institutional Animal Care and Use Committee.

### EAE induction

In order to induce active EAE in mice at the age of 10 weeks, we used a commercial kit (Cat. no. EK-2110, Hooke Laboratories) following its guidelines. Briefly, a subcutaneous injection of an emulsion of myelin oligodendrocyte glycoprotein (MOG_35-55_) in complete Freund’s adjuvant (CFA) was administered followed by two doses of pertussis toxin in phosphate-buffered saline (PBS), each containing 400 ng, which were intraperitoneally administered on the same day as the immunization and the subsequent day. Following immunization, the mice were closely monitored for a period of eight days. At least two researchers conducted daily observations to detect any indications of paralysis or loss of weight. The criteria employed to assess the severity of symptoms were as follows: A score of 0 indicated no clinical signs, 0.5 indicated a partially limp tail, 1 indicated a paralyzed tail, 1.5 indicated a paralyzed tail and one hind limb paresis, 2 indicated uncoordinated movement and hind limb paresis, 2.5 indicated paralysis of one hind limb, 3 indicated complete paralysis of both hind limbs, 3.5 indicated complete paralysis of the hind limb and weakness in forelimbs, 4 indicated complete paralysis of both hind and forelimbs, and 5 indicated that the animal was moribund.

### Immunohistochemical staining

Antigen retrieval for 4 μm sections cut from paraffin-embedded specimens was conducted with citrate buffer (pH 6.0) for 20 min in a pressure cooker (Biocare Medical NxGen). A rabbit monoclonal CD4 antibody (Sino Biological, cat. no. 50134-R001) was used in a 1:4000 dilution with an incubation time of 1 hr in room temperature. BrightVision 1-step detection system goat anti-rabbit HRP (WellMed DPVR110HRP) was used as the secondary antibody, DAB as the chromogen and Mayer’s hematoxylin as the counterstain.

### Histological analyses of the CD4+ positive lymphocytes

CD4 positive T-lymphocytes in the spinal cord were evaluated semi-quantitatively in a scale of 0-3 (0=no positive cells; 1=focal infiltrate less than 50 positive cells; 2= infiltrates with over 50 positive cells; 3= large accumulations of positive cells). The affected levels of spinal cord were also evaluated, ranging from 0-7.

### Cell Isolation

To isolate CNS mononuclear cells, mice with EAE were perfused with ice-cold PBS, and brains and spinal cords were collected and digested in DNase (Sigma-Aldrich) for 30 min at 37°C. Brains and spinal cords were then mechanically dissociated, and mononuclear cells were isolated using Percoll gradient (GE Healthcare). Naive mouse CD4+ T cells were isolated from spleen and lymph node using CD4+ CD62L+ T Cell Isolation Kit, mouse (Miltenyi, cat. no. 130-106-643). Human CD4+ T cells were isolated from human umbilical cord blood using Ficol-Paque PLUS (Cytiva, cat no. 17144003) density gradient centrifugation, followed by purification using CD4+ Dynal positive selection beads (Invitrogen, cat. no. 11331D).

### Cell transfection with siRNA

Splenic mouse CD4+ T cells were transfected using a previously described protocol (*23*). Briefly, cells were activated for 48 h in RPMI media supplemented with 10% FCS, 2 mM l-alanyl-l-glutamine, 1 mM sodium pyruvate, 0.1 mM nonessential amino acids, 55 μM bmercaptoethanol, 100 U/ml penicillin, 100 μg/ml streptomycin, and 10 mM Hepes, in the presence of plate-bound anti-CD3 (5 μg/ml; BD Biosciences, cat. no. 553238) and anti-CD28 (1 μg/ml; BD Biosciences, cat. no. 557393). The preactivated cells were then transfected with siRNAs using an electroporation-based 4D-Nucleofecter system. For each reaction, 2 million cells were resuspended in 15 μl Opti-MEM (Gibco, cat. no. 31985-062) and mixed with 6 μg siRNA in a total volume of 20 μl. The cell-siRNA mixture was transferred to Nucleocuvette Strips and processed using the 4D-Nucleofector 96-well system with program CM137 (Lonza). The following mouse siRNAs were used: siGENOME mouse DZIP3 siRNA SMARTpool (M-051810-01-0010, Dharmacon); siGENOME mouse C330027C09Rik (224171) siRNA SMARTpool (M-055049-01-0010, Dharmacon; CIP2A siRNA), and siGENOME Non-Targeting Pool #2 (D-001206-14-20, Dharmacon).

Human CD4+ T cells were transfected as we have reported earlier (*9*). Briefly, 4 million cells were resuspended in Opti-MEM and mixed with 4 μg siRNA in a total volume of 100 μl. The cell-siRNA mixture was transferred to a cuvette, and nucleofection was performed using the Lonza Nucleofector II with U-14 program (Lonza). The following human siRNAs were used: CIP2A.1 siRNA 5′-CUGUGGUUGUGUUUGCACU-3′ [dT][dT] (*9*); DZIP3 siRNA1 5′-GGGCUCAGCUGGCAAAGUA-3′ [dT][dT], DZIP3 siRNA2 5′-GCCUGGAUGAAUUGCAUAU-3′ [dT][dT] (*24*); Non-targeting siRNA 5′-AAUUCUCCGAACGUGUCACGU-3′ [dT][dT] (*9*).

### Th17 cell culturing

Mouse Th17 cells were generated as previously described (*25*) with slight modifications. In brief, nucleofected cells were activated with plate-bound anti-CD3 (1 μg/ml; BD Biosciences, cat. no. 553238) and anti CD28 (1μg/ml; BD Biosciences, cat. no. 557393) in the presence of IL6 (30 ng/ml; R&D Systems, cat. no. 406-ML), TGFβ1 (5 ng/ml; R&D Systems, cat. no. 240-B), IL7 (5 ng/ml; kindly provided by Associate Professor Alexander Mildner, inFLAMES/University of Turku), and neutralizing antibodies anti-IL4 (10 μg/ml; R&D Systems, cat. no. 559062) and anti-IFNγ (10 μg/ml; R&D Systems, cat. no. 557530). Cell cultures were performed in IMDM media supplemented with 10% fetal calf serum, 2 mM glutamine, 100 iu/mL penicillin, 0.1 mg/mL streptomycin (Sigma, St Louis, MO), and 50 μM β-mercaptoethanol.

Human Th17 cell differentiation was performed as described earlier(*9*). Briefly, CD4+ T cells were activated with plate-bound anti CD3 (3.75 μg/ml; BeckmanCoulter, cat. no. IM1304) and soluble anti-CD28 (1 μg/mL; Beckman Coulter, cat. no. IM1376) antibodies in serum-free X-Vivo 20 medium (Lonza), supplemented with L-glutamine (2 mM, Sigma-Aldrich) and antibiotics (50 U/ml penicillin plus 50 g/ml streptomycin; Sigma-Aldrich), in the presence of IL-6 (20 ng/ml, Roche, cat. no. 11138600 001); IL-1b (10 ng/ml; R&D Systems, cat. no. 201 LB); TGF-b1 (10 ng/ml; R&D Systems Cat. no. 240); anti-IL-4 (1 μg/ml; R&D Systems, cat. no. MAB204), and anti-IFN-g (1 μg/ml; R&D Systems, cat. no. MAB-285).

### Single-cell capture, preparation of sequencing libraries, Single-cell RNA Sequencing and Analysis

Single-cell capture and preparation of sequencing libraries was performed using 10x Genomics Chromium kits and samples were multiplexed using TotalSeq-B antibodies (BioLegend). Before single-cell capture, the cells were suspended in 0.04% BSA in PBS.

For scRNA-seq analysis of CNS cells, single cell partitioning and preparation of sequencing libraries was performed using Chromium Next GEM Single Cell 3’ Reagent Kit v3.1 and the Chromium X instrument, targeting 10,000 cells per sample. The PCR steps were carried out using a Bio-Rad C1000 Touch instrument, using 12 cycles for cDNA pre-amplification and 12 cycles for the sample index PCR. The TotalSeq-B labelled cell pools were processed using 10x Genomics Chromium Single Cell 3′ Library & Gel Bead Kit v3 and the Chromium Controller instrument, targeting 20,000 cells per pool. The PCR steps were carried out using a Veriti cycler (Applied Biosystems/Thermo Fisher), using 11 cycles for cDNA pre-amplification and 12 cycles for the sample index PCR.

For the lymph node samples, the CellPlex-labelled pools were processed using Chromium Next GEM Single Cell 3’ Reagent Kit v3.1 with Feature Barcode technology for Cell Multiplexing and the Chromium X instrument, targeting 30,000 cells per pool. The PCR steps were carried out using a Bio-Rad C1000 Touch instrument, using 12 cycles for cDNA pre-amplification and 13 cycles for the sample index PCR.

The data was analyzed using Seurat (v4.2.0) (*26*) in R (v4.2). For CNS data cells with more than 10% of mitochondrial reads, less than 200 expressed genes or more than 8000 expressed genes were filtered out. For lymph node data cells with more than 10% mitochondrial reads, less than 200 expressed genes or more than 7000 expressed genes were filtered out.

Normalization, identification of highly variable features, scaling, dimensionality reduction with PCA, integration, and clustering were performed per Seurat’s instructions. Uniform manifold approximation and projection (UMAP) was used for data visualization. The clusters were annotated using a reference annotation with SingleR (v1.10.0)(*27*) and manual annotation. Mouse atlas data was used for the reference annotation. For differential expression analysis between KO and WT, cell-level counts were aggregated into sample-level counts in each cell type using muscat (v1.10.1)(*28*). The counts were transformed to logCPM values and ROTS (1.24.0) (*29*) was used for statistical testing of differential expression.

Single-cell suspensions were prepared from lymph node cells of CIP2A^HOZ^ and wild-type mice and subjected to single-cell RNA-seq analysis in two batches. We did not detect any batch effect with the mouse, replicates, or the genotype. We then annotated the clusters based on the expression of marker genes and combined the clusters located in the same UMAP space and with similar gene expression profiles.

### Flow Cytometry

Data were acquired by BD LSR II or BD LSR Fortessa analyzer (BD Biosciences, Franklin Lakes, NJ) at Cell Imaging and Cytometry Core, and analyzed with FlowJo. Following the acquisition of the data, it was analyzed by Flowjo (FlowJo LLC). The following antibodies were used: CD4 PE (BD Biosciences, Cat. No. 553049), CXCR6 APC (BioLegend, Cat. No. 151106), TCRγδ PE (BioLegend, Cat. No. 118107), CD3 BV605 (BD Biosciences, Cat. No. 563004), CD45 V500 (BD Biosciences, Cat. No. 561487), and IL-17 Alexa Fluor 647 (BD Biosciences, Cat. No. 560224).

### TaqMan Quantitative Real-Time-PCR (TaqMan qRT-PCR)

The Qiagen RNeasy® Mini Kit (cat. no. 74106) was used for RNA isolation according to the manufacturer’s instructions. Single-stranded cDNA was synthesized using either SuperScript II Reverse Transcriptase (Invitrogen, cat. no. 18064014) or the SuperScript IV VILO Master Mix with ezDNase (Invitrogen, cat. no. 11766500), following the manufacturer’s protocols. Details of the TaqMan primers and probes are provided in **Table S4**. TaqMan reactions were performed on a QuantStudio 12K Flex Real-Time PCR instrument, and data were analyzed using QuantStudio 12K Flex Real-Time PCR System software (v1.2.3; Thermo Fisher Scientific). Cycle threshold (Ct) values for the internal control genes (EF1α or GAPDH) were subtracted from the Ct values of the target genes to calculate ΔCt (dCt) values.

### ELISA

Secreted IL-17a levels were determined from human Th17 cell culture supernatants at 72 h using the human IL-17a DuoSet ELISA kit (R&DSystems, cat. no. DY317-05, DY008).

### Statistics and Plotting

Statistical significance was determined using two-tailed paired Student’s t-test unless otherwise specified. For EAE scores, statistical significance was assessed using a full linear mixed-effects model in GraphPad Prism, with days post-immunization, genotype, and their interaction included as fixed effects and repeated measurements from individual mice accounted for in the model. The Geisser–Greenhouse correction was applied. Data visualizations were carried out using R, Microsoft Excel and GraphPad Prism.

## AUTHOR CONTRIBUTIONS

M.M.K. designed and performed experiments, analyzed data, prepared figures, and wrote the manuscript. I.S. designed and performed experiments, analyzed data, prepared figures, and contributed to manuscript writing. S.J. and J.S. analyzed scRNA-seq data. E.R. maintained the mouse colony and contributed to experiment planning, data analysis, and data interpretation.

R.B. performed experiments and processed scRNA-seq samples. M.H.K. designed and performed experiments, analyzed data, and prepared figures. R.K. performed experiments and analyzed data. M.M.B. and U.R. contributed to manuscript writing. J.K. and E.Y. contributed to EAE experiments and provided technical expertise. P.P.H. and L.S. provided critical feedback and insight for EAE model and contributed to experimental design and data interpretation. T.L. contributed to scRNA-seq data analysis and interpretation. L.L.E. supervised J.S. and S.J. and provided expertise in computational data analysis. J.W. provided scientific input, expertise, mice, and reagents. O.R. designed experiments, analyzed data, and contributed to writing the manuscript. R.L. designed and supervised the study and wrote the manuscript. U.U.K. designed experiments, analyzed data, prepared figures, and wrote the manuscript. All authors reviewed and edited the manuscript and approved the final version.

## DATA AVAILABILITY

The scRNA-seq data from lymph node and CNS will be deposited at GEO.

## ETHICAL APPROVAL

All procedures and protocols regarding animal experiments were approved by the National Project Authorization Board of Finland (project license ESAVI/24048/2021, renewed under decision ESAVI/25753/2025) and conducted in accordance with the EU Directive 2010/EU/63 and Finnish national legislation (497/2013 and 564/2013) on the protection of animals used for scientific purposes.

Umbilical cord blood was obtained from healthy neonates, both sexes, at Turku University Central Hospital for this study. The usage of the cord blood of unknown donors was approved by the Ethics Committee of Hospital District of Southwest Finland (Dated: 24.11.1998; ethical approval number: 323). As the subjects were neonates, informed consent was obtained from the parents or their legal representatives. Cord blood samples have been collected anonymously without any information on the sex of the neonates.

## ACKNOWLEDGEMENTS

We thank Marjo Hakkarainen and Sarita Heinonen for their exceptional technical assistance. We thank the staff of Turku University Hospital, Department of Obstetrics and Gynecology, Maternity Ward, for the cord blood collection. We would also like to acknowledge the invaluable support provided by the Turku Bioscience Centre’s core facilities, namely Finnish Functional Genomics Centre, Single Cell Core and Cell Imaging Core Facility and the Turku Center for Disease Modeling, which are all supported by Biocenter Finland. Personnel of the Central Animal Laboratory are acknowledged for their expert animal care and technical support. We also thank the Finnish Centre for Scientific Computing for providing efficient servers and data analysis resources.

MMK was supported by the University of Turku Graduate School Turku Doctoral Programme of Molecular Medicine (TuDMM) and a central grant from the Finnish Cultural Foundation supported. RL was supported by the Research Council of Finland (RCF) Centre of Excellence in Molecular Systems Immunology and Physiology Research (2012-2017) grant 250114; and by other RCF grants 292335, 294337, 292482, 31444. RL was also supported by grants from the JDRF, the Sigrid Jusélius Foundation and the Finnish Cancer Foundation the Novo Nordisk Foundation grant NNF19OC0057218, Jane and Aatos Erkko Foundation grant. The generation of the mouse model by JW laboratory was supported by the Sigrid Jusélius Foundation and the Finnish Cancer Foundation. LLE received support from the European Research Council (ERC) grant number 677943 and the European Union’s Horizon 2020 research and innovation program grant number 955321. LLE was also supported by the RCF grants 310561, 314443, 329278, 335434, 335611, and 341342. LLE were supported by the Sigrid Jusélius Foundation. RL and LLE were supported by Turku Graduate School University of Turku, Åbo Akademi University, InFLAMES Flagship Programme of the RCF under grant number 337530, Biocenter Finland, and ELIXIR Finland.

