## Supplementary figures and tables for "Genetic Disruption at the CIP2A Locus Modulates T Cell Responses and Attenuates Experimental Autoimmune Encephalomyelitis"

**Figure S1**

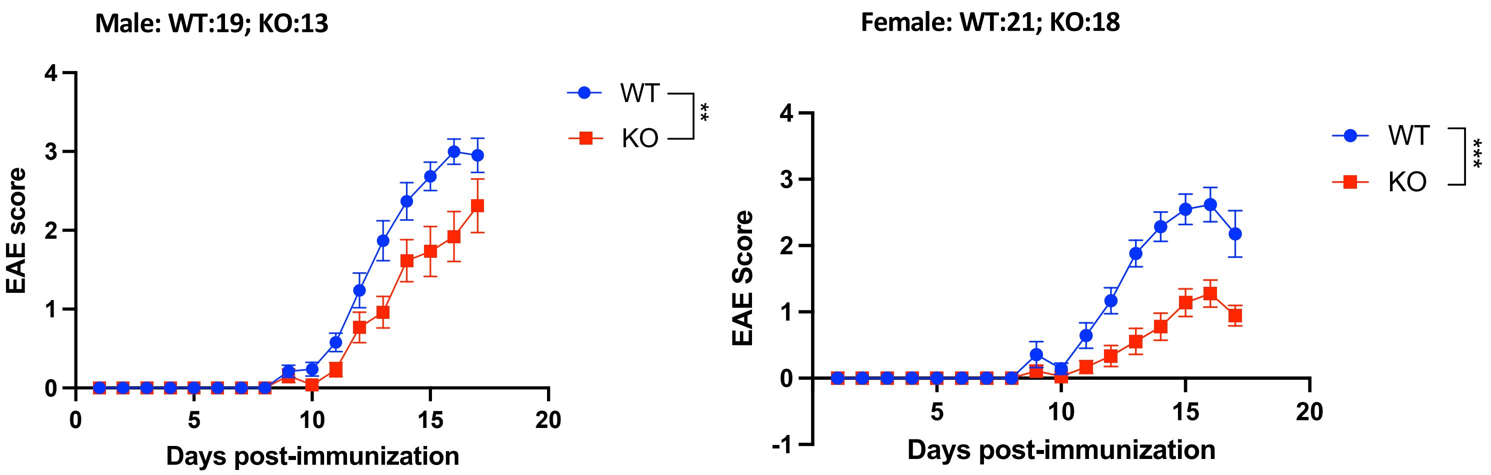


**Figure S1. Genetic disruption at the Cip2a locus ameliorates EAE in female and male mice.** Clinical EAE scores in male (left) and female (right mice). Number of WT and KO mice are shown above the figures. Statistical significance was assessed using a full linear mixed-effects model in GraphPad Prism, with days post-immunization, genotype, and their interaction included as fixed effects and repeated measurements from individual mice accounted for in the model. The Geisser–Greenhouse correction was applied. ** and *** denote P value <0.001 and <0.0001, respectively.

### Figure S2

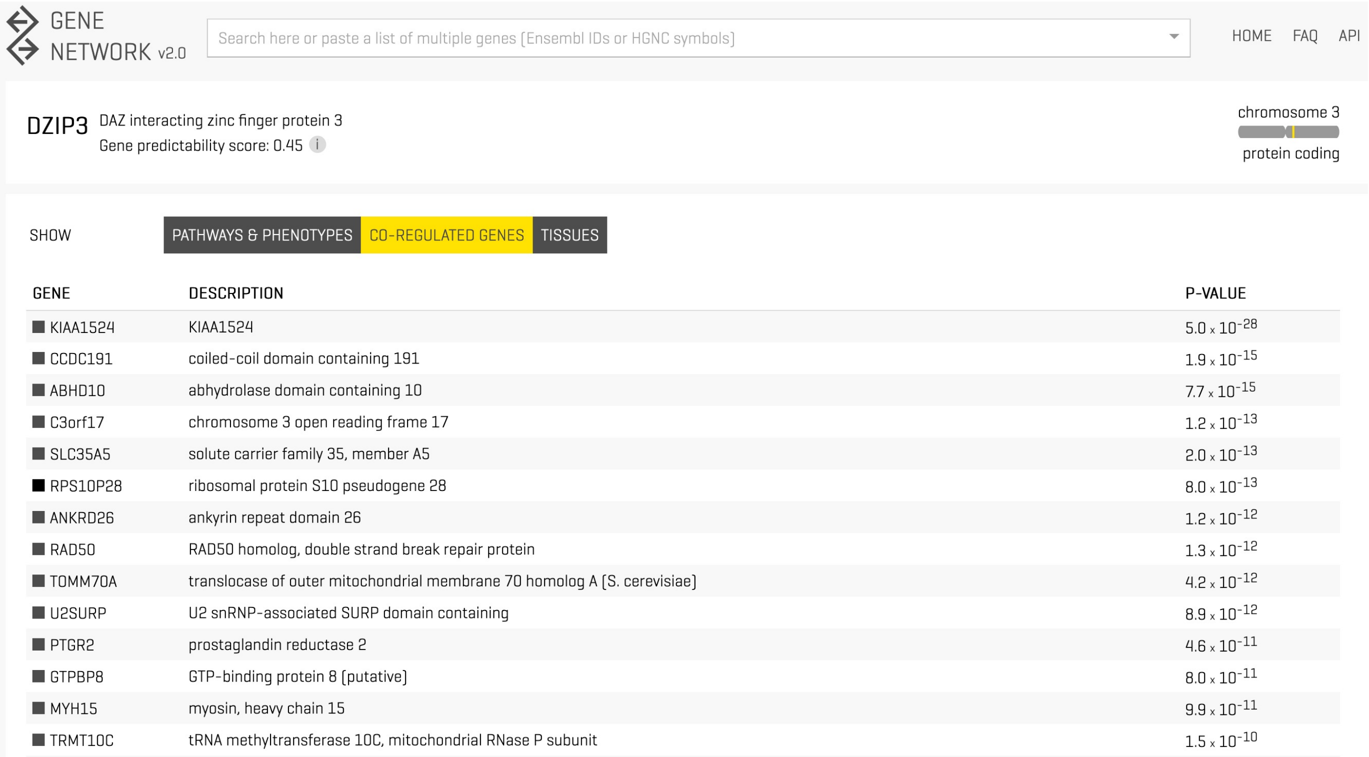

GENE NETWORK v2.0

Search here or paste a list of multiple genes (Ensembl IDs or HGNC symbols)

HOME FAQ API

DZIP3 DAZ interacting zinc finger protein 3

Gene predictability score: 0.45

chromosome 3

protein coding

SHOW PATHWAYS & PHENOTYPES CO-REGULATED GENES TISSUES

| GENE | DESCRIPTION | P-VALUE |
|---|---|---|
| KIAA1524 | KIAA1524 | $5.0 \times 10^{-28}$ |
| CCDC191 | coiled-coil domain containing 191 | $1.9 \times 10^{-15}$ |
| ABHD10 | abhydrolase domain containing 10 | $7.7 \times 10^{-15}$ |
| C3orf17 | chromosome 3 open reading frame 17 | $1.2 \times 10^{-13}$ |
| SLC35A5 | solute carrier family 35, member A5 | $2.0 \times 10^{-13}$ |
| RPS10P28 | ribosomal protein S10 pseudogene 28 | $8.0 \times 10^{-13}$ |
| ANKRD26 | ankyrin repeat domain 26 | $1.2 \times 10^{-12}$ |
| RAD50 | RAD50 homolog, double strand break repair protein | $1.3 \times 10^{-12}$ |
| TOMM70A | translocase of outer mitochondrial membrane 70 homolog A (S. cerevisiae) | $4.2 \times 10^{-12}$ |
| U2SURP | U2 snRNP-associated SURP domain containing | $8.9 \times 10^{-12}$ |
| PTGR2 | prostaglandin reductase 2 | $4.6 \times 10^{-11}$ |
| GTPBP8 | GTP-binding protein 8 (putative) | $8.0 \times 10^{-11}$ |
| MYH15 | myosin, heavy chain 15 | $9.9 \times 10^{-11}$ |
| TRMT10C | tRNA methyltransferase 10C, mitochondrial RNase P subunit | $1.5 \times 10^{-10}$ |

**Figure S2. CIP2A is the top coregulated gene with DZIP3.** The figure shows the most significant genes whose expression coregulated with that of DZIP3. The co-expression analysis is based on thousands of publicly available RNA-seq data as analyzed online at genenetwork.nl.

**Table S1**: Differentially abundant cell types in the scRNA-seq data from KO vs WT CNS samples

| | logFC | logCPM | F | PValue | FDR |
|---|---|---|---|---|---|
| Gamma Delta T cells | 1.455504087 | 12.85318769 | 4.587119227 | 0.055544782 | 0.995658777 |
| DCs | 0.795665974 | 14.17341454 | 2.594283252 | 0.135673357 | 0.995658777 |
| Flt3+ DCs | 1.031465623 | 14.07237587 | 1.592919584 | 0.233824437 | 0.995658777 |
| Fcnb+ monocyte progenators | 0.623169049 | 14.10268686 | 0.558960018 | 0.470851184 | 0.995658777 |
| Hexb+ Microglia | -0.409694137 | 16.19378353 | 0.529705331 | 0.482002867 | 0.995658777 |
| B cells | 0.370434829 | 14.06173118 | 0.462934179 | 0.510384996 | 0.995658777 |
| NK cells | 0.33030403 | 13.56315727 | 0.360166535 | 0.560627601 | 0.995658777 |
| Ms4a7+ Monocytes | 0.545679032 | 16.07646532 | 0.350613226 | 0.566113441 | 0.995658777 |
| Proliferating CD8 T cells | 0.373039326 | 13.10704915 | 0.325518735 | 0.579828288 | 0.995658777 |
| Neutrophils | -0.211446975 | 18.91029084 | 0.258391524 | 0.621310895 | 0.995658777 |
| Ccr2+ monocytes | 0.291669169 | 14.4166627 | 0.185209096 | 0.675281702 | 0.995658777 |
| Hexb+ Hspa1a+ Hspa1b+ Microglia | 0.166694456 | 15.67630672 | 0.09932464 | 0.758565728 | 0.995658777 |
| CD4 Effector Cells | 0.099544701 | 16.3902169 | 0.037334835 | 0.85032202 | 0.995658777 |
| Tissue resident memory CD8 T cells | -0.119725508 | 13.30111587 | 0.031468582 | 0.862437647 | 0.995658777 |
| Foxp3+ Tregs | -0.018533928 | 13.22948543 | 0.001682204 | 0.968022427 | 0.995658777 |
| Erythroid cells | -0.027806873 | 15.1406095 | 0.000877668 | 0.976913128 | 0.995658777 |
| Icos low CD 8 T cells | 0.004890337 | 14.84726097 | 6.07E-05 | 0.993925782 | 0.995658777 |
| Ccr7 high Naive CD4 T cells | 0.004296172 | 13.77964938 | 3.10E-05 | 0.995658777 | 0.995658777 |

**Table S2**: Differentially expressed genes in different clausters in the scRNA-seq data from KO vs WT CNS samples

| logFC | pvalue | padj | gene | cluster |
|---|---|---|---|---|
| -4.2703063 | 2.85E-05 | 0 | Stfa1 | Hexb+ Microglia |
| -3.6048743 | 1.27E-05 | 0 | Cip2a | Hexb+ Microglia |
| -2.6402782 | 8.36E-05 | 0 | Fzd1 | Neutrophils |
| -5.1823573 | 2.32E-05 | 0 | Csta2 | Neutrophils |
| -4.8250368 | 1.17E-05 | 0 | Stfa1 | Neutrophils |
| -3.3817271 | 3.49E-05 | 0 | Gm17103 | Neutrophils |
| -6.0862866 | 1.48E-06 | 0 | Arhgap31 | Neutrophils |
| -4.2898717 | 2.47E-05 | 0 | Ccdc80 | Neutrophils |
| -4.1789093 | 1.50E-05 | 0 | Dzip3 | Neutrophils |
| -3.8134871 | 2.00E-05 | 0 | Cip2a | Neutrophils |
| -4.3202743 | 2.17E-05 | 0 | Stfa1 | CD4 Effector Cells |
| -3.1272007 | 0.00010928 | 0 | Gm17103 | CD4 Effector Cells |
| -3.0499801 | 2.80E-05 | 0 | Plcxd2 | CD4 Effector Cells |
| -2.963879 | 2.49E-05 | 0 | Dzip3 | CD4 Effector Cells |
| -3.1622077 | 1.60E-05 | 0 | Cip2a | CD4 Effector Cells |
| 4.49199217 | 0.00016531 | 0 | Sdc4 | Erythroid cells |
| 3.56214641 | 0.00042249 | 0 | Fam43a | Erythroid cells |
| -5.7285747 | 3.10E-05 | 0 | Stfa1 | Erythroid cells |
| -5.3636777 | 5.02E-05 | 0 | Cip2a | Erythroid cells |
| 2.68055003 | 8.48E-06 | 0 | Gbp8 | DCs |
| -3.0212476 | 4.14E-05 | 0 | Dzip3 | DCs |
| -5.8121777 | 1.69E-06 | 0 | Cip2a | DCs |
| -7.9615371 | 2.13E-05 | 0 | Jchain | B cells |
| -9.1155834 | 1.26E-05 | 0 | Igha | B cells |
| -3.9261411 | 0.00013296 | 0 | Iglv1 | B cells |
| -3.5499345 | 7.57E-05 | 0 | Plcxd2 | B cells |
| 1.65489632 | 1.61E-05 | 0 | Rpgr | NK cells |
| -4.2789125 | 3.40E-05 | 0 | Stfa1 | Fcnb+ monocyte progenators |
| -5.7375346 | 2.03E-06 | 0 | Cip2a | Fcnb+ monocyte progenators |
| -4.3773127 | 0.00014876 | 0 | Il17a | Proliferating CD8 T cells |
| -3.4561187 | 2.32E-05 | 0 | Cip2a | Proliferating CD8 T cells |
| 1.71453351 | 4.91E-05 | 0 | St5 | Ttr+ CNS cells |
| -2.7965043 | 9.18E-06 | 0 | Dzip3 | Ms4a7+ Monocytes |

**Table S3**: Differentially abundant cell types in the scRNA-seq data from KO vs WT lymph node samples

| logFC | pvalue | padj | gene | cluster |
|---|---|---|---|---|
| -1.2189604 | 2.57E-05 | 0 | Pde3b | AICDA+ B cells |
| -1.6249218 | 3.13E-05 | 0 | Slc15a3 | AICDA+ B cells |
| -1.5944718 | 5.23E-06 | 0 | Tbc1d15 | AICDA+ B cells |
| 1.23100255 | 2.06E-05 | 0 | Cpped1 | Anxa1+ basophils |
| -1.4744148 | 1.66E-05 | 0 | Gtf2f2 | Anxa1+ basophils |
| 1.43286693 | 4.31E-05 | 0 | Hira | Anxa1+ basophils |
| -2.8103632 | 8.98E-07 | 0 | Arhgap31 | B cells |
| 1.9335605 | 3.56E-05 | 0 | B4galt4 | B cells |
| 1.64310693 | 3.88E-05 | 0 | Ccdc38 | B cells |
| -2.0821781 | 7.52E-06 | 0 | Cip2a | B cells |
| -3.047881 | 4.49E-07 | 0 | Dzip3 | B cells |
| 2.02772435 | 2.19E-06 | 0 | Gm38393 | B cells |
| 2.5129444 | 1.74E-06 | 0 | Hsd3b2 | B cells |
| 2.24456233 | 2.64E-06 | 0 | Nr4a2 | B cells |
| -1.7458143 | 1.11E-05 | 0 | Plcxd2 | B cells |
| -1.5236023 | 3.75E-05 | 0 | Ell | Ccl22+ monocytes |
| -0.9989624 | 5.70E-05 | 0 | Gpatch3 | CCND1+ mDCs |
| 1.04167617 | 2.56E-05 | 0 | Mocs1 | CCND1+ mDCs |
| -1.1791471 | 4.49E-05 | 0 | BC052040 | CD4_cl32 |
| -1.3126086 | 1.02E-05 | 0 | Igsf9b | CD44 high Treg |
| -0.8854661 | 1.50E-05 | 0 | Zfp235 | CD44 high Treg |
| -1.4799152 | 1.56E-05 | 0 | Tchh | Clec4a3+ mDcs |
| -1.9473652 | 3.91E-06 | 0 | Fbxo46 | Gamma Delta T cells I |
| -1.2612028 | 2.59E-05 | 0 | Klhl42 | Gamma Delta T cells I |
| -1.6006939 | 4.92E-06 | 0 | Atp9b | Il18+ myeloid cells |
| -1.8003488 | 5.55E-06 | 0 | F10 | Il18+ myeloid cells |
| -1.8509593 | 5.43E-05 | 0 | Metap1 | Il18+ myeloid cells |
| -3.4523291 | 1.06E-05 | 0 | Arhgap31 | IL1B+ monocytes |
| -2.7702221 | 3.91E-05 | 0 | Csta2 | IL1B+ monocytes |
| 3.7918497 | 6.33E-06 | 0 | Cstdc5 | IL1B+ monocytes |
| -4.7600986 | 2.55E-06 | 0 | Stfa1 | IL1B+ monocytes |
| -2.7086905 | 2.08E-05 | 0 | Cip2a | Plasma cells |
| -2.5774261 | 1.77E-05 | 0 | Dzip3 | Plasma cells |
| -3.4010578 | 1.40E-05 | 0 | Ighv1-74 | Plasma cells |
| -4.5432997 | 3.66E-05 | 0 | Ighv3-6 | Plasma cells |
| -0.9206516 | 1.57E-06 | 0 | Gm48194 | Tfh |
| -1.7216994 | 4.67E-05 | 0 | Znrf3 | Tfh |
| -1.0537481 | 3.84E-05 | 0 | Nutf2-ps1 | Th17 |

**Table S4.** TaqMan qRT-PCR primers and probes

| Gene name | mouse | human |
|---|---|---|
| CIP2A | Probe: Universal probe library probe # 19 (Roche Life Science) F-primer: 5' -CTGCGTTGAACATTGAGGAC- 3' R-primer: 5' -ATCTCATAAACATCCATGATGTCAG- 3' | Probe: Universal probe library probe # 69 F-primer: 5' -GAACAGATAAGAAAAGAGTTGAGCATT- 3' R-primer: 5' -CGACCTTCTAATTGTGCCTTTT- 3' |
| DZIP3 | TaqMan Gene Expression Assay, Assay ID # mm01343818_m1 (Thermo Fisher Scientific) | TaqMan Gene Expression Assay, Assay ID # Hs00978125_m1 |
| IL17A | Probe: Universal probe library probe # 34 F-primer: 5' -GATTTTCAGCAAGGAATGTGG- 3' R-primer: 5' -CATTGTGGAGGGCAGACAAT- 3' | Probe: Universal probe library probe #8 F-primer: 5' -TGGGAAGACCTCATTGGTGT- 3' R-primer: 5' -CGGATTTCGTGGGATTGTGAT- 3' |
| IL17F | none | Probe: Universal probe library probe #10 F-primer: 5' -GGCATCATCAATGAAAACCA- 3' R-primer: 5' -TGGGGTCCCAAGTGACAG- 3' |
| CCR6 | none | Probe: 5' 6-FAM/-AGGCGACTAAGTCATTCCGGCTCCG/TAMRA/- 3' F-primer: 5' -GACCGGTACATCGCCATTGTA- 3' R-primer: 5' -TCGTGCGCGGTAGTGTTCT- 3' |
| mEF1-α | Probe: 5' 6-FAM/-CTTCAAATTCACCAACACCAGCAGCAAC/TAMRA/- 3' F-primer: 5' -AGGCTGACTGTGCTGTCCTGAT- 3' R-primer: 5' -GCCCGTTCTTGGAGATACCA- 3' | Probe: 5'/56-FAM/AGCGCCGGCTATGCCCCTG/3BHQ_1/- 3' F-primer: 5' -GCCGTGTGGCAATCCAAT- 3' R-primer: 5' -CTGAACCATCCAGGCCAAAT- 3' |
| GAPDH | TaqMan Gene Expression Assay, Assay ID # mm99999915_g1 | none |
